# Expected and observed genotype complexity in prokaryotes: correlation between 16S-rRNA phylogeny and protein domain content

**DOI:** 10.1101/494625

**Authors:** Jasper J. Koehorst, Edoardo Saccenti, Vitor Martins dos Santos, Maria Suarez-Diez, Peter J. Schaap

## Abstract

**Background:** The omnipresent 16S ribosomal RNA gene (16S-rRNA) is commonly used to identify and classify bacteria though it does not take into account the distinctive functional characteristics of taxa. We explored functional domain landscapes of over 5700 complete bacterial genomes, representing a wide coverage of the bacterial tree of life, and investigated to what extent the observed protein domain diversity correlates with the expected evolutionary diversity, using 16S-rRNA as metric for evolutionary distance.

**Results:** Analysis of protein domains showed that 83% of the bacterial genes code for at least one of the 9722 domain classes identified. By comparing clade specific and global persistence scores, candidate horizontal gene transfer and signifying domains could be identified. 16S-rRNA and functional domain content distances were used to evaluate and compare species divergence and overall a sigmoid curve is observed. Already at close 16S-rRNA evolutionary distances, high levels of functional diversity can be observed. At a larger 16S-rRNA distance, functional differences accumulate at a relatively lower pace.

**Conclusions:** Analysis of 16S-rRNA sequences in the same taxa suggests that, in many cases, additional means of classification are required to obtain reliable phylogenetic relationships. Whole genome protein domain class phylogenies correlate with, and complement 16S-rRNA sequence-based phylogenies. Moreover, domain-based phylogenies can be constructed over large evolutionary distances and provide an in-depth insight of the functional diversity within and among species and enables large scale functional comparisons. The increased granularity obtained paves way for new applications to better predict the relationships between genotype, physiology and ecology.

## Introduction

The most commonly used method to classify bacteria and to identify new isolates is the direct comparison of the omnipresent 16S ribosomal RNA (16S-rRNA) gene sequence^1,2^ with highly curated 16S-rRNA gene sequence databases^3–8^.

Using only the 16S-RNA gene for taxonomic characterisations presents limitations and disadvantages. First, arbitrary minimal sequence similarity thresholds are used as working boundaries for differentiating between taxonomic ranks. Although these thresholds prove to be very useful for classification purposes, they are subject to progressive insights and are limited as there is no biological meaning attached to it^9^. For instance, the minimal sequence similarity threshold for species delineation, proposed for the 16S-rRNA gene, has changed over time from 97% to 98.7%10,11 and even at this updated stringency level, the resolution is too limited for a definite species classification of some phylogenetic groups,^12^. Second, a restriction to the analysis of sequence variations in a single gene does not take into account the distinctive functional characteristics of the different prokaryotic taxa nor can it explain the genotypic, and the consequently phenotypic, differentiation observed between strains due to events such as gene loss or acquisition.

Alternative, inter-genomic BlastN-based sequence similarity methods exist that take into account full genome sequences. Examples are Average Nucleotide Identity (ANI)^13,14^, Genome Blast Distance Phylogeny (GBDP)^15^ or a combination of 16S-rRNA sequence similarity and ANI values^16^. These methods help to increase taxonomic coherence at the smaller evolutionary distances, but are less suitable to monitor the impact of mutation and (strain specific) gene loss and horizontal gene transfer (HGT).

To better understand the impact of gene loss and HGT and to improve the characterisation of functional diversity, the analysis needs to be performed beyond genome sequence similarity comparison by considering protein function. Protein encoding genes reveal a modular design, with domains forming distinct globular structural and functional units. Bacterial innovation is in part driven by gain, loss, duplication and rearrangement of these functional units, resulting in the emergence of proteins with new domain combinations^17,18^. Thus, a direct comparison of protein domain content should be able to reconstruct bacterial phylogeny independent of gene sequence similarity^19^ and as such may serve as a better indicator of shared physiology and ecology^20,21^.

In this study we present an exhaustive exploration of the functional landscape of over 5700 complete bacterial genomes representing a wide coverage of the bacterial tree of life and investigated to what extent protein domain diversity correlates with taxonomic diversity using the 16S-rRNA gene sequence as metrics for evolutionary distances.

## Results

We analysed 5713 fully sequenced publicly available, bacterial genomes corresponding to a wide range of different bacterial lineages (57 classes, 243 families, 818 genera and multiple strains of 1330 species), providing a good representation of the bacterial diversity observed in nature (See supplementary file S1 for more information). Genome sizes varied from 0.1 Mbp up to 13 Mbp. To avoid technical bias due to the use of different annotation strategies, all genomes were *de-novo* re-annotated with SAPP^22^ (see Methods section for details). The total number of genes varied from 167 *(Candidatus Tremblaya princeps)* to 9968 *(Streptomyces bingchenggensis* BCW-1).

### 16S-rRNA variability within and between species

From the 5713 completely sequenced genomes, 25098 complete 16S-rRNA genes could be retrieved. On average the predicted length of the 16S-rRNA gene was 1531 ± 94 nt (See supplementary Figure Supplementary file S1) and 84% of the completed genomes (4772) contained between two and fifteen copies of the 16S-rRNA gene (Figure 1). The 16S-rRNA genes from phylogenetic cohesive groups of at least 50 strains were further analysed at family level. As can be seen in Figure 1B, among different families there is a diverse variation in copy number. As already has been observed^23^, while in some families the 16S-rRNA copy number is largely restricted to a single copy gene, copy number in others ranged from 1 to 15 copies. Furthermore, 52% of the analysed genomes contained two or more non-identical copies of the 16S-rRNA gene. Intragenomic sequence variation reflected an overall sequence identity of 99.6 (+0.4 / −2)%, which is higher than the currently accepted 98.7% threshold for species delineation.

**Figure 1.**
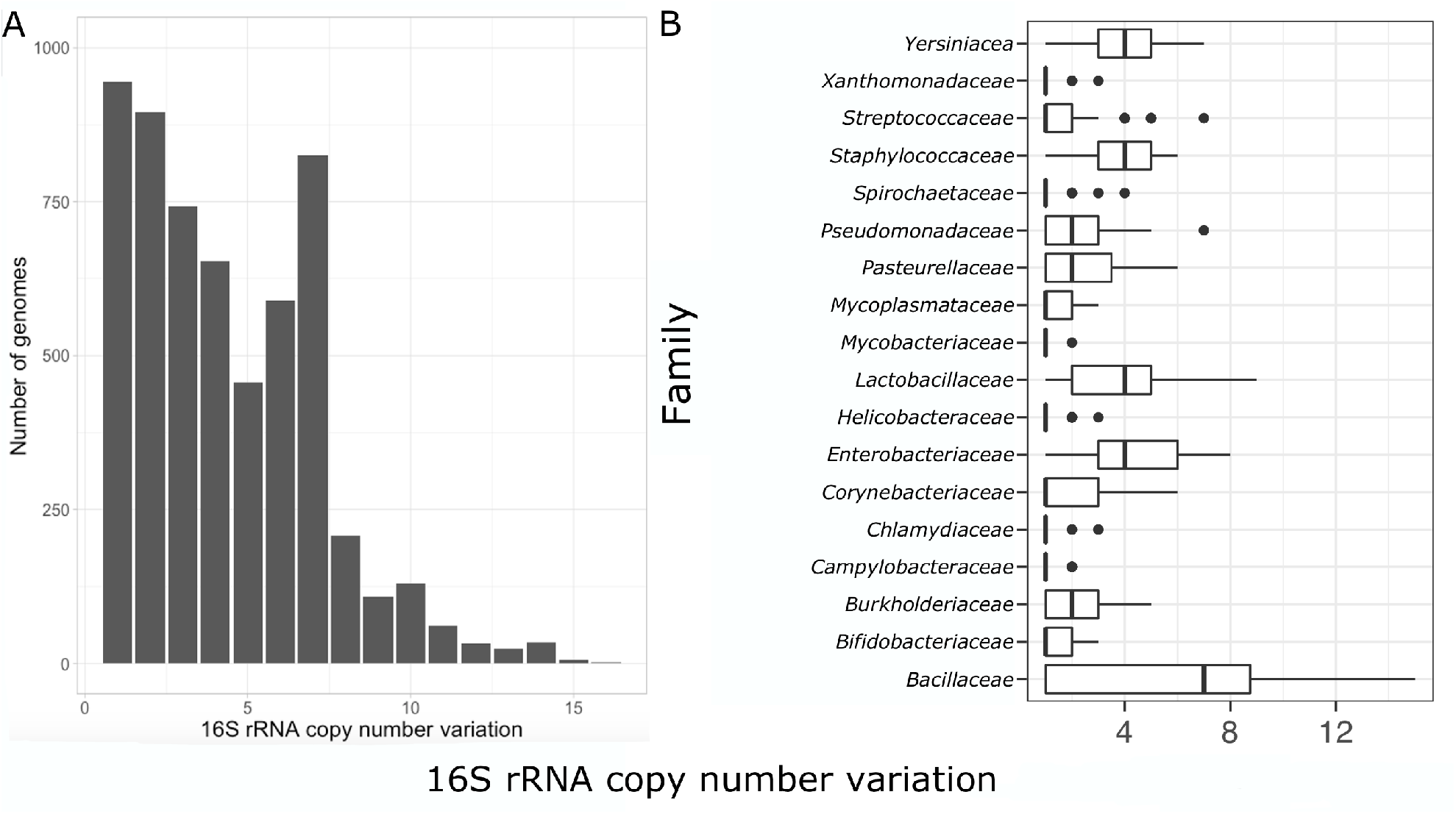
16S-rRNA copy number variation. A) 16S-rRNA gene copy number variation in the complete set B) Copy number variation at family level; families represented by more than 50 strains were analysed.

For the complete set of genomes, a species network based on pair-wise 16S-rRNA sequence similarity scores was built. In this network, nodes represent genomes and edges were drawn between nodes when the 16S-rRNA showed at least 98.7% identity. Network connectivity analysis identified 2025 connected components (subnetworks). For further study, 294 subnetworks linking ten or more nodes, were were selected. In thirty-two of these subnetworks, taxonomic inconsistencies were observed as they linked genomes assigned to two or more species. The majority (30) of these taxonomic inconsistent subnetworks linked species belonging to the same genus. However, two subnetworks were identified that linked species from different genera. The first contained species of the *Escherichia* and *Shigella* genera. The second subnetwork showed even more diversity and contained members of the *Citrobacter, Enterobacter, Klebsiella, Kosakonia, Raoultella* and *Salmonella* genera (Figure 2). Both subnetworks eventually belong to the *Enterbacteriaceae* family. Overall, network analysis suggested that in many cases the 98,7% identity threshold is not sufficient and additional means of classification are required to obtain reliable phylogenetic relationships.

**Figure 2.**
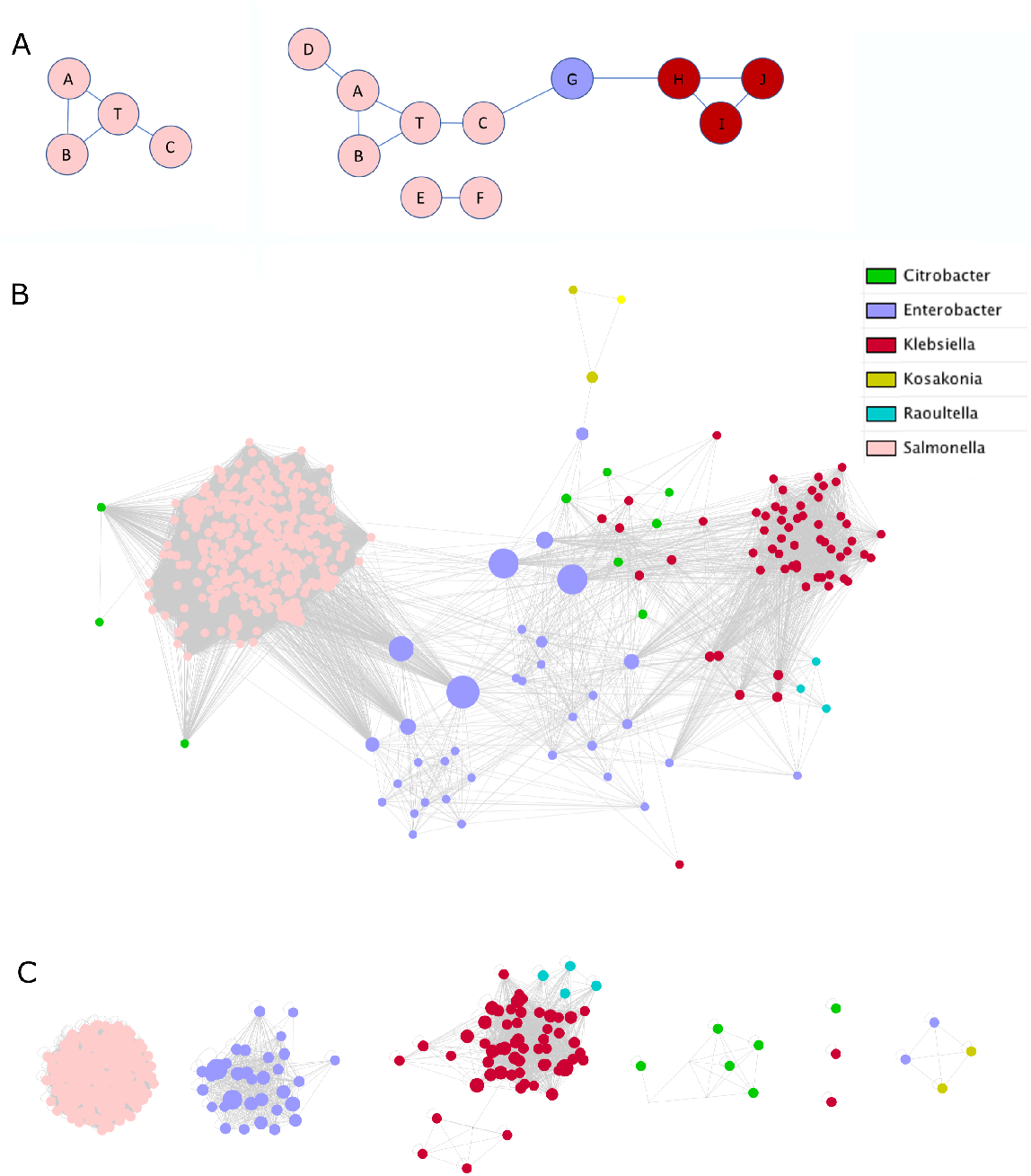
Topology of the similarity subnetwork of *Enterbacteriaceae*. Nodes represent genomes and edges are drawn if the 16S-rRNA identity >98.7%. A) Network topologies with colours indicating the different species groups. Left panel, unambiguous species assignment; strains A,B and C are directly connected to type strain T. Right panel: Observed topology. Leave node strain D is in the cluster but has no direct link with type strain T. 16S-rRNA sequences of strain E and strain F are below the set similarity threshold and form an unlinked subnetwork. Strain G of the blue species functions as an articulation point linking the pink and red species subnetworks. B) Subnetwork linking six different genera based on the 16S-rRNA gene sequences using a sequence similarity threshold >98.7%. Size of each node is dependent on the betweenness centrality. *Enterobacter* is the main component that connects the different genera as no direct linkage between *Salmonella* and *Klebsiella* is observed. Three strains of *Citrobacter* have a direct connection to *Salmonella* and are disconnected from other *Citrobacter* strains. One *Enterobacter* (*Enterobacter sp. R4-368*) is isolated from the rest and is only connected to *Kosakonia*. The *Raoultella* genera have a close similarity to some of the *Klebsiella* strains. C) Topology of domain-class content subnetworks of the same strains using as threshold a binary distance ≤0.1. Distinct subnetworks are observed. *Salmonella* is now completely separated from the other genera; *Enterobacter, Klebsiella* and *Citrobacter* also form distinct clusters with a few members forming separate subnetworks.

### Protein domain architectures

By breaking proteins into domains and using precomputed profile hidden Markov models (pHMM) to classify these domains, a semantically consistent classification of encoded protein functions can be obtained^20^. As a pHMM gives greater weight to matches at conserved sites they are also better for remote homology detection than standard sequence similarity-based methods^24^. To obtain such protein classification the 18949996 inferred protein sequences were scanned for the presence of Pfam domains^25^. A total of 15747648 protein sequences were found to contain at least one domain instance (83.1%) and in total 9722 distinct protein domain classes were detected (See supplementary file S1 for more details). Two Pfam domains were discovered in 17.7% (3345544) and three or more domains in 6.4% (1205997) of these proteins (Table 1). Thus, the majority of the bacterial proteins appear to be single domain proteins (Figure 3A). Moreover, we observed that most multiple domain proteins appear to contain domain repetitions. Similar domain distributions were obtained when individual genomes were analysed, indicating that this is a general property of the architecture of bacterial genomes (Figure 3B).

**Figure 3.**
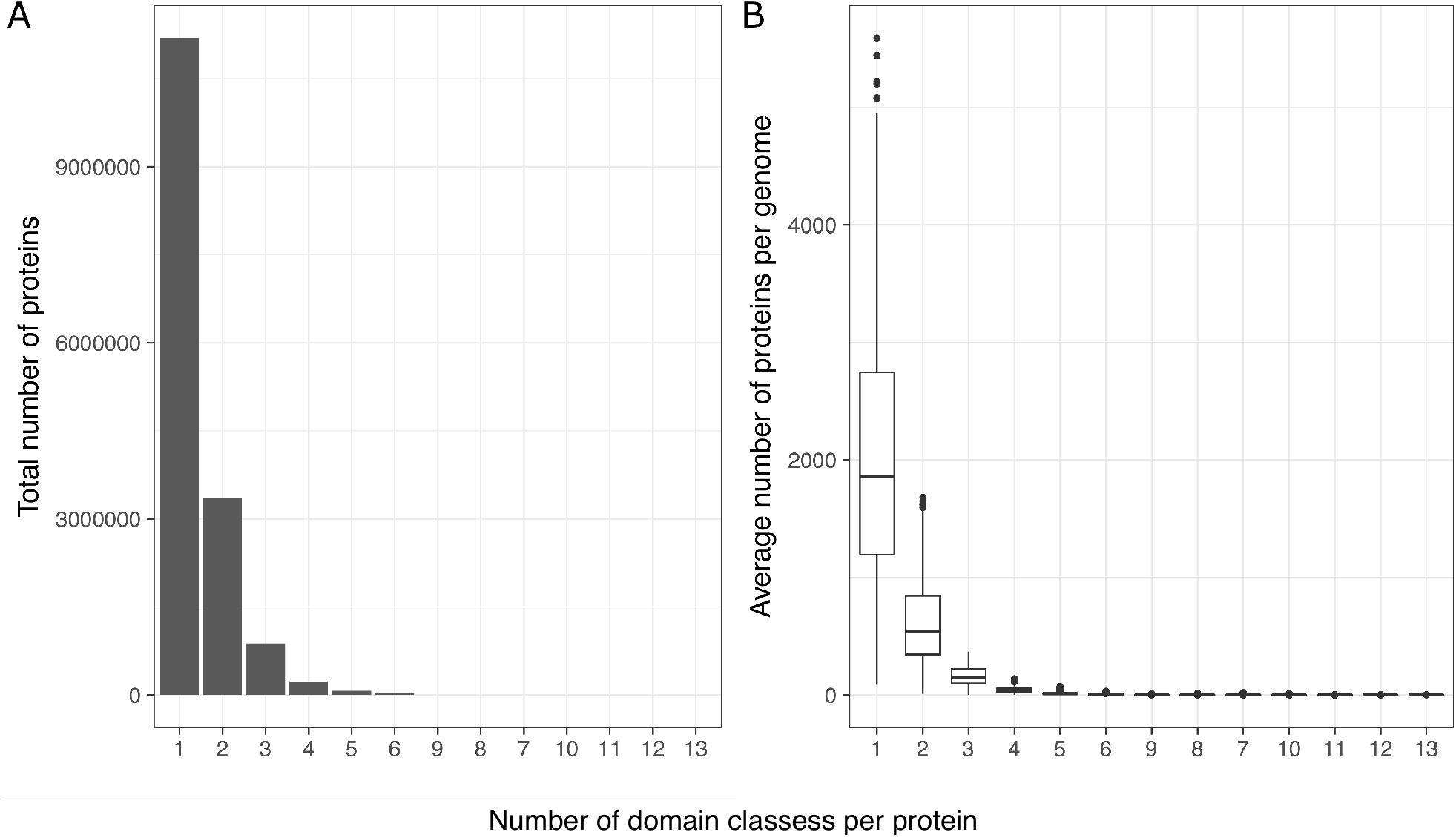
Frequency distribution of 1,2,…,13 domain classes. A) In the full data set. B) In each genome

**Table 1.** Overview of the number of proteins and corresponding protein domain content. The majority of the proteins (83.1%) contained at least a single domain and only a few (1.74%) contained more than 3 domains.

|  | Number of proteins | Fraction |
| --- | --- | --- |
| **All proteins** | 18949996 | 100% |
| **>0 domains** | 15747648 | 83.1% |
| **1 domain** | 11196108 | 59.1% |
| **2 domains** | 3345544 | 17.7% |
| **3 domains** | 875863 | 4.6% |
| **>3 domains** | 330133 | 1.74% |
| **>10 domains** | 15457 | 0.08% |
| **>50 domains** | 208 | 0.0011% |

### Genome distribution of protein domains

The distribution of the domain classes across the studied genomes is shown in Figure 4. Panel A shows that there is a direct correlation between the genome size and the total number of domains detected. A non-linear relationship is observed between the total number of protein domains and the total number of protein domain classes indicating that domain copy numbers, but not so much the number of domain classes, increase in the larger genomes (Figure 4 panel B). On average, we counted 2.02 domain copies per genome. This copy number, however, showed a large variability, ranging from 1.07 copies for *Carsonella ruddii* (strain PV)^26^ to 4.58 copies for *Streptomyces bingchenggensis* (strain BCW-1)^27^.

**Figure 4.**
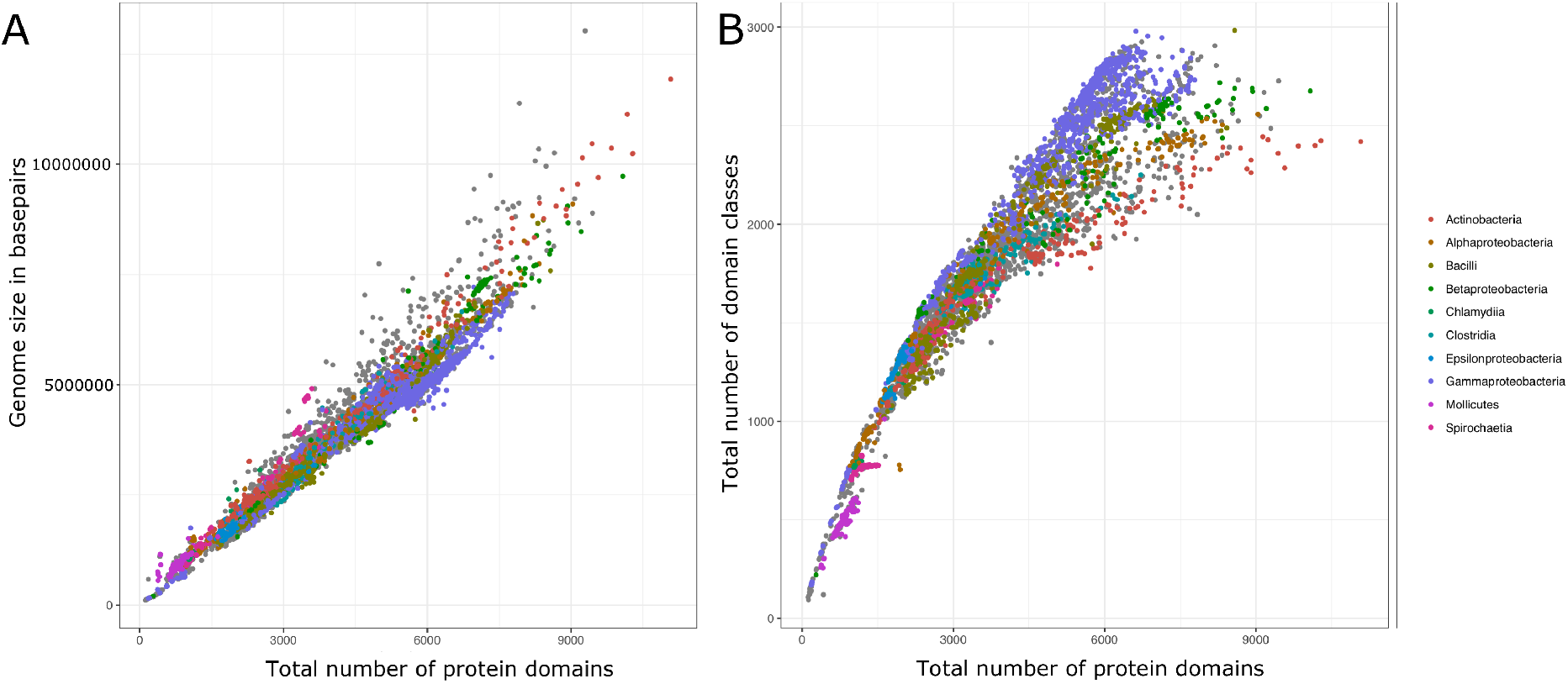
Distribution of the domain classes across bacterial genomes. A) Correlation between genome size and number of protein domains. B) Correlation between the total number of domains and total number of domain classes. A non-linear relation is observed, suggesting that in the larger genomes an increase in domain copy number is favoured over an increase in domain classes.

### Domain persistence and analysis of the pan- and core-domainomes

In total, 9722 domain classes were detected. The overall persistence (the fraction of the genomes sharing a given domain class) is shown in Figure 5. Only 324 domain classes were ubiquitous in over 95% of the analysed genomes. Three domains, PF00009, (GTP-binding elongation factor family), PF01479, (S4 domain) and PF03144 (Elongation factor Tu domain 2) were shown to persist in all genomes. Additionally, a small number of domains were found to be present in over 99.9% of the studied genomes, PF00012 (Hsp70 protein), PF00318 (Ribosomal protein S2), PF00380 (Ribosomal protein S9/S16), PF00679 (Elongation factor G C-terminus), PF01926 (50S ribosome-binding GTPase), PF02811 (PHP domain), PF07733 (Bacterial DNA polymerase III alpha subunit) and PF14492 (Elongation Factor G, domain II). Among the studied genomes there are domain classes with a high copy number. The domain with the highest copy number is PF00005, representing the ATP-binding domain of ABC transporters, with on average 62.9 copies per genome, yet the domain is absent in twelve small-sized genomes.

**Figure 5.**
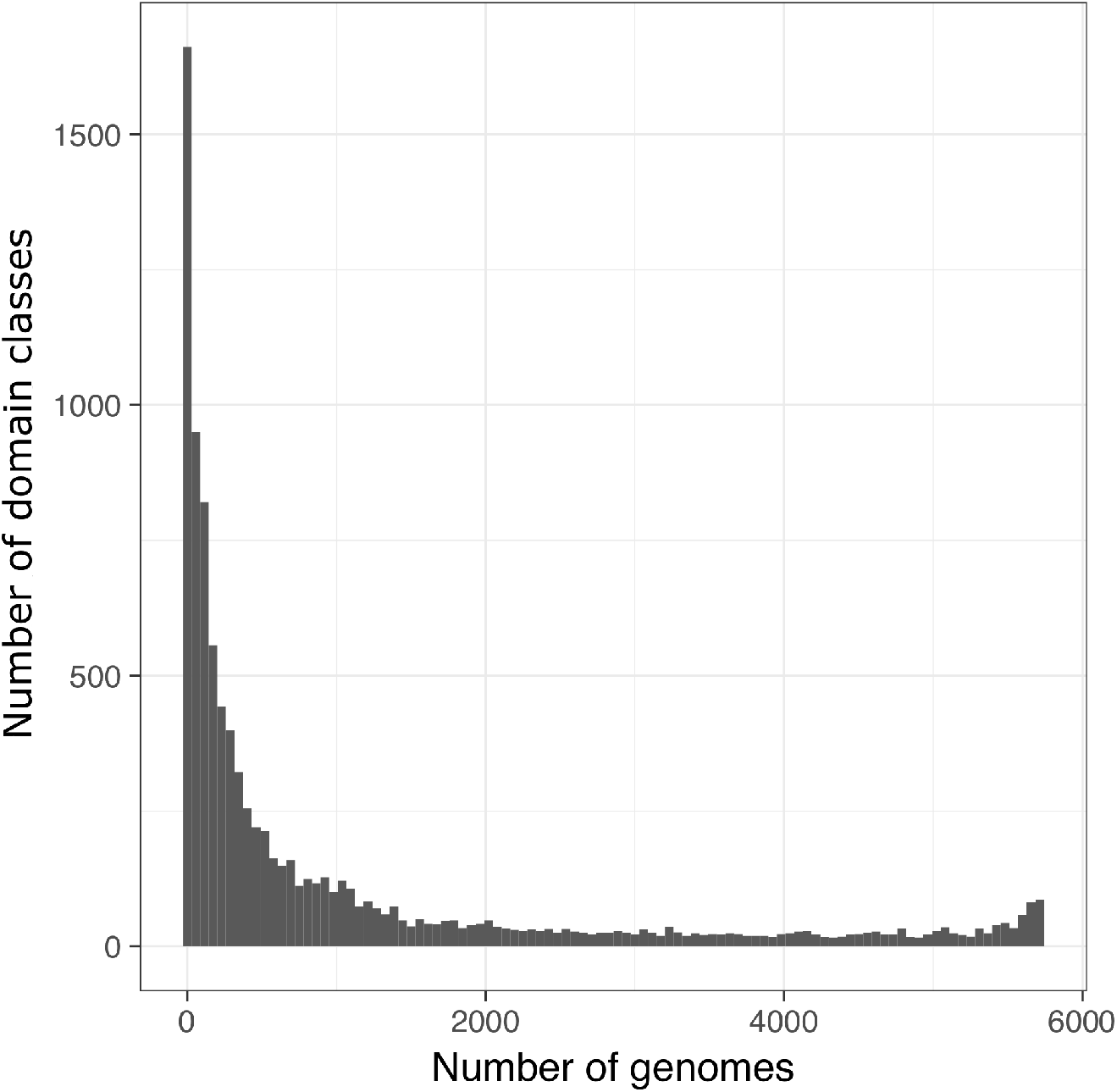
Distribution of domain classes over 5713 genomes

Accurate measurements of the pan- and core-domainome sizes would entail knowledge of the functional content of every single organism in the corresponding group. We have estimated their respective sizes for the 18 families that contained more than 50 members each (Figure 6A). The largest observed pan-domainome was of *Bacillaceae* with 4783 protein domain classes. The largest core was observed for *Yersiniaceae* (1844 domain classes) (Figure 6B).

**Figure 6.**
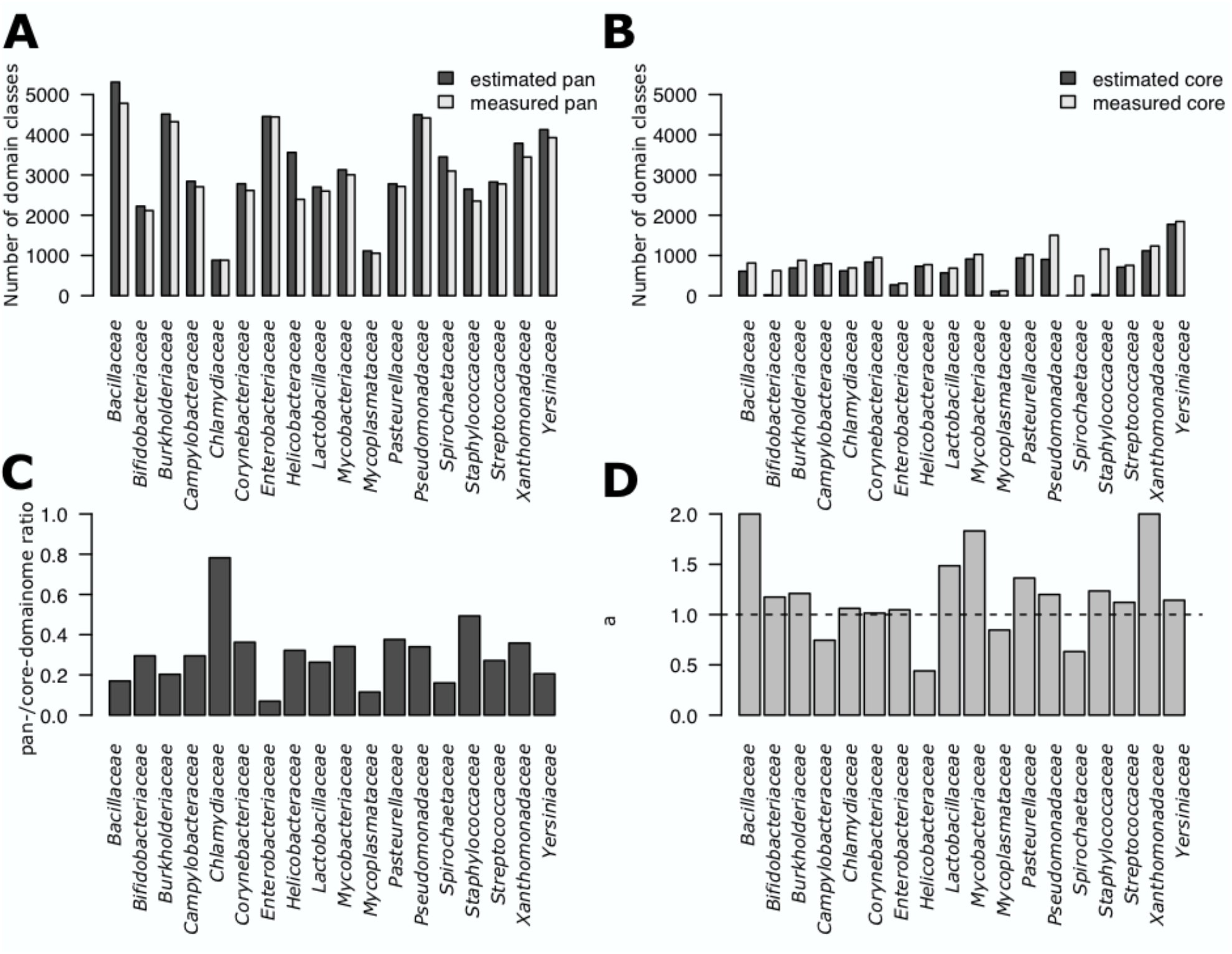
Persistence analysis of families with more than 50 members. The estimated pan-domainome (Panel A) and estimated core (Panel B) shows a large degree of variability ranging from 78% for *Chlamydiaceae* and 7% for Enterobacteriaceae. The conservation ratio of the pan/core (Panel C) shows that in only *Chlamydiaceae* more than half of the protein domain content is conserved. The family pan-genome is closed (Panel D) when *α*>1.

When analysing the genomes of the *Chlamydiaceae* family, 78% of the protein domain classes are conserved. In contrast, the core of *Enterobacteriaceae* only covers 7% of the in total 4444 domains (Figure 6C). This is mostly due to the size of the genomes from the *Moranella, Riesia, Blochmannia* and *Ishikawaella*^28^, genera as they are smaller than 1 Mbp, encoding as low as 444 genes, whereas the average genome size of *Enterobacteriaceae* is 4.8 Mbp, encoding on average 4510 genes. When excluding the small sized genomes, the core increases to 938 protein domains with a slightly smaller pan-domainome of 4441 yielding a 21% ratio between the core and pan-domainome. This shows the impact of including or excluding specific genomes in the analysis, as a single or few genomes can reduce the core significantly, thereby possibly eluting important information.

Openness of the pan-domainome provides another indication of the relative impact of horizontal acquisition and vertical transmission in shaping the domainome. Fitting a Heap’s law, we estimated whether the pan-domainome for each of the largest families was either open or closed by fitting the decay parameter of a Heap’s law function, *α*. The pan-domainome is closed when *α* >1.0 and open when *α* < 1.0. The majority of the bacterial families here considered showed a closed pan-domainome (Figure 6D). For *Enterobacteriaceae* the Heap’s parameter dropped from *α*=1.21 to *α*=1.17 upon removal of the previously indicated smaller genomes.

### Signifying domains and horizontal domain transfer

Log persistence scores (log-P) were calculated for each of the domain classes present in the pan-domainomes from the five most abundant monophyletic species groups *(Chlamydia trachomatis* (74), *Escherichia coli* (105), *Helicobacter pylori* (65), *Salmonella choleraesuis* (350) and *Staphylococcus aureus* (74).) As null-model we consider the persistence of the domain in the full set of 5713 genome sequences.

For a small set of domain classes high (log-P) scores were obtained and are likely signifying domain classes (Table 2, Figure 7 and Supplementary Table S3 logP). On the other end of this scale we find a large amount of domain classes with negative log-P scores. These incidental domains have a low to very low intra-species persistence which suggests that they may have been acquired by horizontal gene transfer. Unlike the high scoring domains most of them have been assigned a molecular, often metabolic, function.

**Figure 7.**
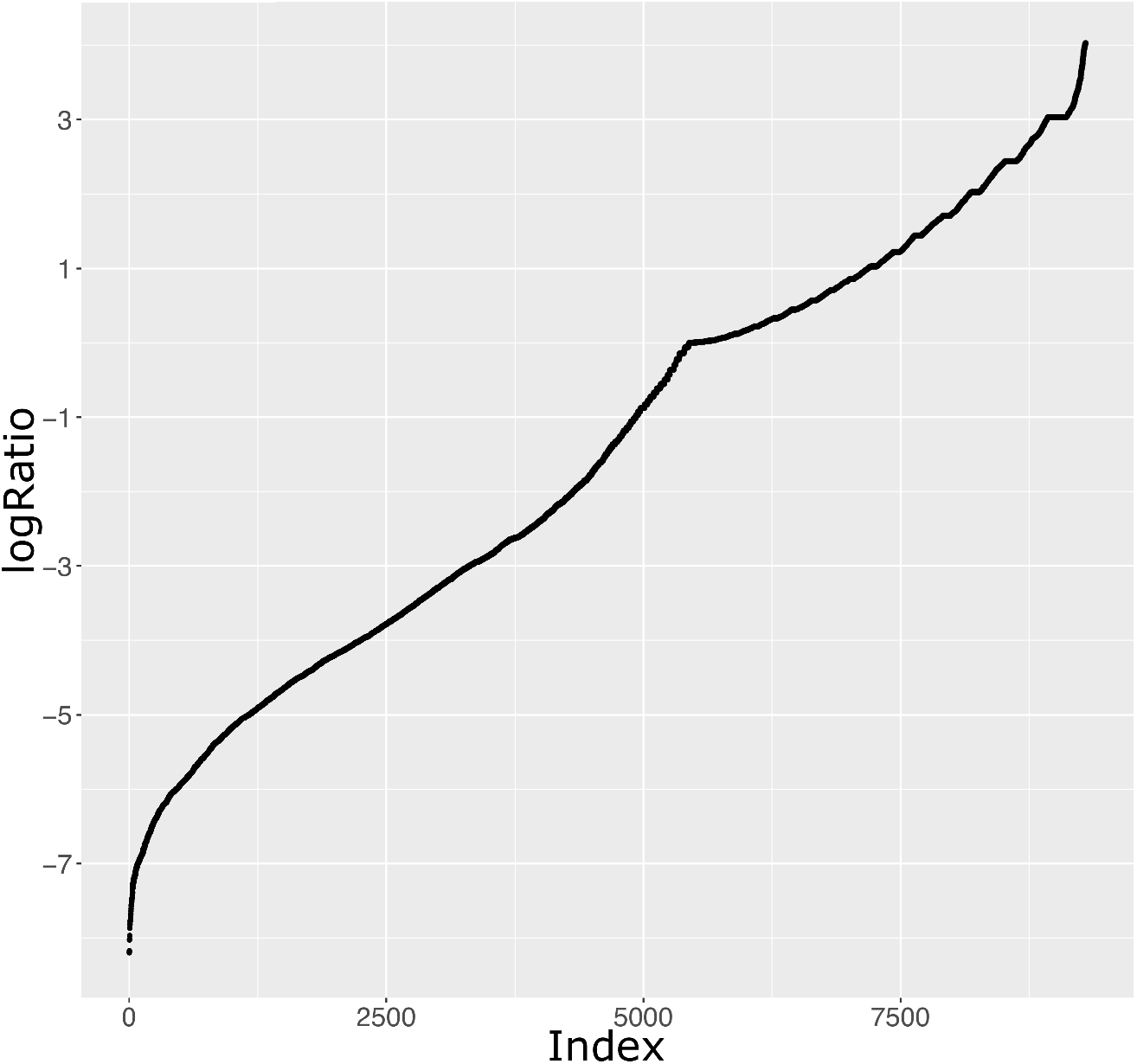
Persistence scores of *Salmonella choleraesuis* protein domain classes. For each domain class present in the *S. choleraesuis* pan-domainome, persistence scores are compared with the pan-domainome persistence scores obtained from the complete set of 5713 genomes.

**Table 2.** *Salmonella choleraesuis* top 25 signifying and incidental domains

| PFAM | log-P | Global persistence | Description |
| --- | --- | --- | --- |
| PF09460 | 4.025 | 0.056 | Saf-pilin pilus formation protein |
| PF06767 | 4.017 | 0.062 | Sif protein |
| PF05364 | 4.013 | 0.062 | Salmonella type III secretion SopE effector N-terminus |
| PF07108 | 4.013 | 0.062 | PipA protein |
| PF16583 | 4.012 | 0.061 | Zinc-regulated secreted antivirulence protein C-terminal domain |
| PF16728 | 3.991 | 0.061 | Domain of unknown function (DUF5066) |
| PF15942 | 3.976 | 0.064 | Domain of unknown function (DUF4751) |
| PF07824 | 3.969 | 0.064 | Type III secretion chaperone domain |
| PF09052 | 3.965 | 0.064 | Salmonella invasion protein A |
| PF05775 | 3.950 | 0.059 | Enterobacteria AfaD invasin protein |
| PF11047 | 3.941 | 0.065 | Salmonella outer protein D |
| PF05925 | 3.914 | 0.066 | Enterobacterial virulence protein IpgD |
| PF08052 | 3.906 | 0.066 | PyrBI operon leader peptide |
| PF13998 | 3.873 | 0.068 | MgrB protein |
| PF02510 | 3.858 | 0.069 | Surface presentation of antigens protein |
| PF13979 | 3.852 | 0.068 | SopA-like catalytic domain |
| PF02090 | 3.840 | 0.070 | Salmonella surface presentation of antigen gene type M protein |
| PF04741 | 3.815 | 0.071 | InvH outer membrane lipoprotein |
| PF07487 | 3.811 | 0.071 | SopE GEF domain |
| PF09119 | 3.801 | 0.072 | SicP binding |
| PF05688 | 3.794 | 0.071 | Salmonella repeat of unknown function (DUF824) |
| PF03433 | 3.759 | 0.074 | EspA-like secreted protein |
| PF09599 | 3.759 | 0.074 | Salmonella-Shigella invasin protein C (IpaC_SipC) |
| PF10940 | 3.737 | 0.074 | Protein of unknown function (DUF2618) |
| PF05689 | 3.727 | 0.074 | Salmonella repeat of unknown function (DUF823) |
| ... |  |  |  |
| PF13442 | -7.502 | 0.518 | Cytochrome C oxidase, cbb3-type, subunit III |
| PF09424 | -7.522 | 0.525 | Yqey-like protein |
| PF01769 | -7.533 | 0.529 | Divalent cation transporter |
| PF06750 | -7.546 | 0.534 | Bacterial Peptidase A24 N-terminal domain |
| PF09084 | -7.555 | 0.537 | NMT1/THI5 like |
| PF03309 | -7.557 | 0.538 | Type III pantothenate kinase |
| PF06271 | -7.573 | 0.544 | RDD family |
| PF12802 | -7.610 | 0.558 | MarR family |
| PF01628 | -7.642 | 0.571 | HrcA protein C terminal domain |
| PF01593 | -7.680 | 0.586 | Flavin containing amine oxidoreductase |
| PF10397 | -7.689 | 0.589 | Adenylosuccinate lyase C-terminus |
| PF00355 | -7.732 | 0.607 | [2Fe-2S] domain |
| PF01220 | -7.743 | 0.612 | Dehydroquinase class II |
| PF14693 | -7.769 | 0.623 | Ribosomal protein TL5, C-terminal domain |
| PF01809 | -7.769 | 0.623 | Haemolytic domain |
| PF02616 | -7.771 | 0.624 | Segregation and condensation protein ScpA |
| PF04079 | -7.781 | 0.629 | Segregation and condensation complex subunit ScpB |
| PF03448 | -7.815 | 0.644 | MgtE intracellular N domain |
| PF07521 | -7.823 | 0.647 | Zn-dependent metallo-hydrolase RNA specificity domain |
| PF01883 | -7.868 | 0.668 | Iron-sulfur cluster assembly protein |
| PF02686 | -7.969 | 0.716 | Glu-tRNA<sup>Gln</sup> amidotransferase C subunit |
| PF02637 | -8.019 | 0.741 | GatB domain |
| PF02934 | -8.026 | 0.744 | GatB/GatE catalytic domain |
| PF01425 | -8.173 | 0.824 | Amidase |
| PF00825 | -8.196 | 0.838 | Ribonuclease P |
| | | | Number of strains analysed 350; $\alpha=0.89$ |

### Co-evolution of bacterial 16S-rRNA and whole genome domain content

Protein domains provide a formal description of genome encoded functionalities, each contributing to bacterial genotypic complexity. The functional relatedness of an arbitrary pair of genomes can thus be determined by finding the fraction of encoding domain classes in common relative to the the number of domain classes present in each of these genomes. Through inclusion of the 16S-rRNA data the co-evolution of bacterial 16S-rRNA gene sequences with genotypic complexity can be studied (Figure 8). In panel A the distribution of domain based distances is plotted using a binary dissimilarity score. Likewise in panel D, the distribution of 16S-rRNA sequence distances is plotted. Panel C shows a pairwise comparison between 16S-rRNA distances and functional distances for the analysed genomes. Finally, panel B, presents a schematic representation of the relationship between the two methods.

**Figure 8.**
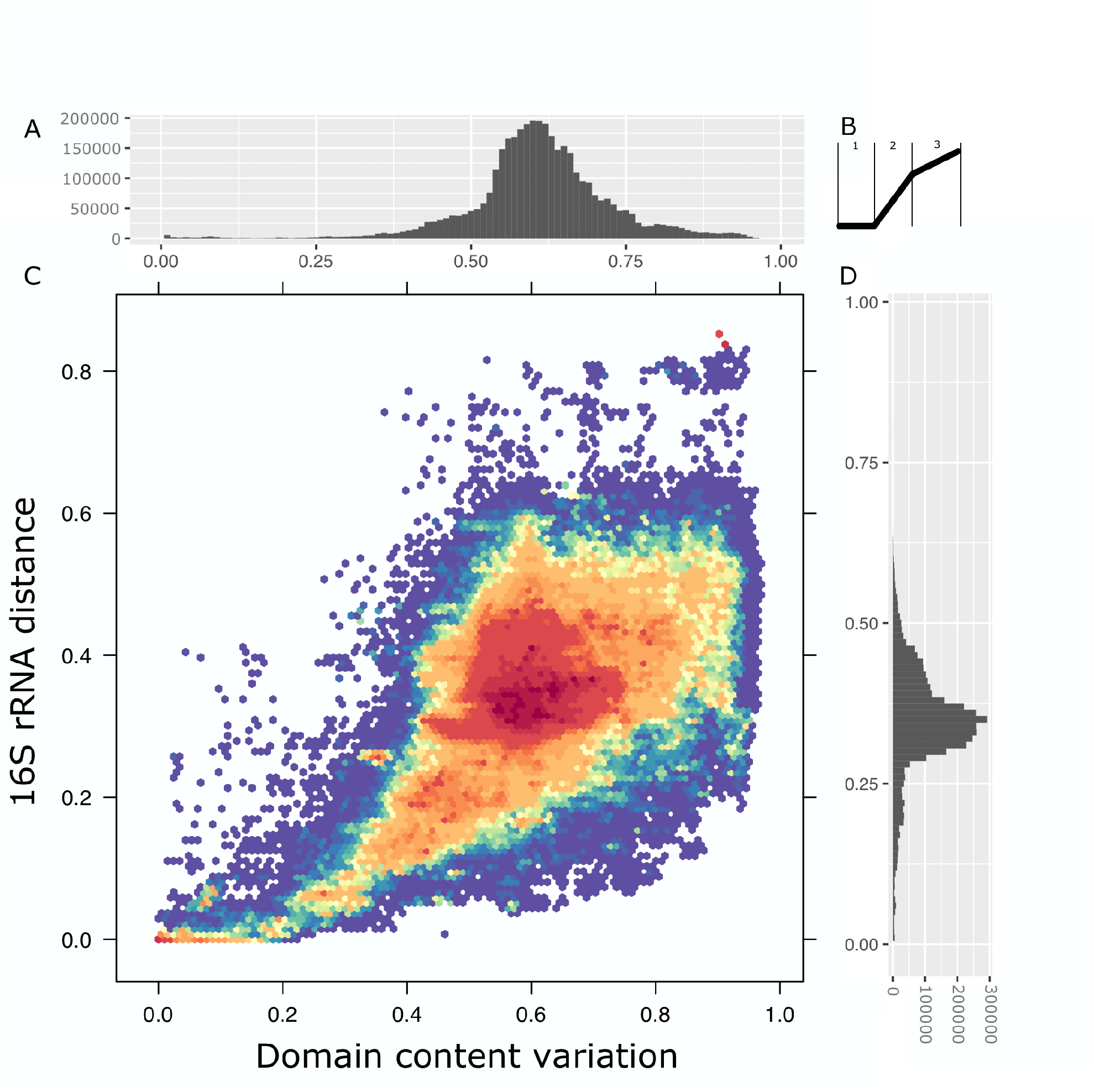
Distance comparison of the 16S-rRNA gene with the functional diversity. A) Distribution of domain based distances. B) Schematic representation of the three stages of diversification. 1) a fast-short-term evolution, as evolutionary distances measured by 16S-rRNA remain small, while functional diversification has already taken place. 2) long-term evolution, in which functional diversification occurs at a scale compatible with diversification by 16S-rRNA sequence evolution. 3) The distance of the 16S-rRNA remains behind the functional diversity as the 16S-rRNA distance can only diverse so far without loss of function. C) Comparison between pairwise 16S-rRNA distances and pairwise functional distances. Color indicates density of points, blue and red indicate lower and higher density respectively D) Distribution of 16S-rRNA based distances.

Overall, a good agreement is found between both approaches to evaluate species divergence. Analysis of the 16S-rRNA distances shows a marked differentiation in the [0.3, 0.35] interval, which appears as a steep increase in the abundance of instances of these distance values (Figure 8D). These differentiations correspond to lineage boundaries (specifically class and phylum differences). This increased density corresponds to the higher density in the center of the plot (Figure 8C), that reflects that most of the performed comparisons involve members distantly related in the evolutionary scale. This is also apparent on the higher number of instances of functional differences in the [0.6, 0.7] interval (Figure 8A), however functional differences accumulate more gradually, and no steep increase is observed.

The relationship between the two methods to evaluate species differences can be approximated through a sigmoidal curve and three regimes can be distinguished (Figure 8B). Species at close evolutionary distances show a broad range of functional similarity (Figure 8B region 1). A high diversity is observed, so that genomes with high similarity regarding their 16S-rRNA can show high functional diversity. The second region shown in Figure 8B, region 2, corresponds to regions of relatively large genetic differentiation (class differences) that accumulate functional differences at a relatively lower pace. Finally, the third region (region 3) corresponds to very distant species that as expected, have a large degree of functional differentiation.

In addition to functional similarities between evolutionary close strains, Figure 8C also indicates the presence of functionally very similar but evolutionary distant genomes. These are to be found in the region with low domain content variation (<0.05) and a large 16S-rRNA distance (>0.4). *Gluconacetobacter diazotrophicus* PAl 5, *Moraxella catarrhalis* BBH18 and *Pseudomonas aeruginosa* 39016 are some examples. Similar results are obtained when the analysis is repeated considering all available genomes. The presence of more than one copy of the 16S-rRNA gene may introduce a larger variability, however the overall agreement of 16S-rRNA classification remains the same.

## Discussion

For several decades 16S-rRNA sequence similarity scores provided a good working metric for prokaryotic taxonomic classifications, but because of the ever-expanding sequence databases and the increased taxonomic complexity the limitations of this approach are emerging. Here, we have used a set of 5713 complete genomes to evaluate the predictive power of pair-wise 16S-rRNA sequence similarity scores on the diversity and taxonomic classification of these genomes.

We observed intragenomic variation of 16S-rRNA gene sequences, but further analysis showed that within the selection, this variation is limited and well above the currently advised species threshold of 98.7%, meaning that regardless of the selected copy, the same taxonomic classification should be obtained.

A network approach was subsequently used to study pair-wise 16S-rRNA sequence similarities between the 5713 sequenced strains (Figure 2). By using the currently accepted 98.7% minimal sequence similarity threshold, optimally this approach should lead to 1330 separate species networks, each containing all sequenced strains of a defined species and each individual node within such subnetwork should at least have a direct link to the node that represents the reference or type strain (Figure 2 panel A). However, many more subnetworks were obtained and what was observed is that strains of the same species are in separate subnetworks. Additionally, strains with intermediate 16S-rRNA sequences were present functioning as articulation points merging what should have been independent species subnetworks. (Figure 2 panel B and C). With the continuous addition of new 16S-rRNA sequences it is likely that species amalgamation will become more frequent. In the light of this, a more appropriate approach would be to consider the similarity threshold as a confidence level. In this way, there is a high probability that two sequences with a 16S-rRNA sequence identity below the selected threshold belong to different species. This provides a probabilistic interpretation to the threshold.

We used Pfam protein domain-class content to study strain diversity. Protein domains are considered to be distinct functional units and as such responsible for a particular function or interaction. The Pfam 30 protein family database consists of 16306 domain families or classes^29^ of which 9721 were present in the studied dataset. Furthermore, we found that approximately 83% of the protein-encoding genes harbour at least one Pfam domain suggesting that the encoded domain-class content may provide a good metric to study strain diversity.

The core-genome of a taxonomic group contains genes that are present in all members of that group whereas the pan-genome contains all the different genes that can be found in any member of the population^30^. Here we extended the idea to protein domain classes, as has been previously reported^21,31^. We observed that most domain classes have a low persistence overall (Figure 5), but as shown in Figure 6, by adding taxonomic information, distinct sets of domain classes accumulate in the core domainomes of the various clades, suggesting that these core sets are somehow contributing to the physiology and ecology of these clades.

At family level, the pan to core domainome ratio is observed to be on average below 0.4 (Figure 6), but at lower taxonomic ranks this ratio increases. For *C. trachomatis* this ratio was determined to be 0.96, for *Escherichia coli* 0.58, for *Helicobacter pylori 0.83* and for *Staphylococcus aureus* 0.76. We assumed that species core domainomes would consist of signifying or even species-specific domain classes and domain-classes representing essential metabolic functions. We expected that signifying domain-classes are only highly persistent within a clade but that domain-classes representing metabolic functions would be widely spread. For each domain class present in the pan-domainome of five selected species we calculated the ratio between clade specific persistence and global persistence (log-P scores) using a null-model that assumes that domain-classes are evenly distributed over the strains. The analysed species contributed to 6.2% or less of the total number of strains.

Top log-P scoring domains mostly corresponded to domains of unknown function (DUF) or domains involved in signal transduction whereas, being omnipresent, metabolic functions were underrepresented. Of the 25 top scoring domains, 6 in *Salmonella choleraesuis*, 15 in *Chlamydia trachomatis*, 8 in *Escherichia coli*, 3 in *Helicobacter pylori* and 11 in *Staphylococcus aureus* corresponded to a DUF class. For the *Mycoplasma* species it has been established that many DUFs are essential for growth^32,33^ and at least four of the DUFs in the present study, two specific for *Escherichia coli* (PF07041 and PF10897) and two for *Helicobacter pylori* (PF12033 and PF10398) indeed have been characterised as being essential^34^. Between these five species, top scoring domains also show no significant overlap suggesting that they are evolutionary conserved and may have a prominent role in shaping the species. Protein domain classes with the lowest persistence ratio’s are likely HGT candidates. Functionally, most of them represent a metabolic function suggesting, as has been reported^35,36^, that horizontal gene transfer is an important source of metabolic diversity.

The impact of the presence of these signifying domains in the core domainome is demonstrated in Figure 2C. Nodes from the *Enterobacteriaceae* subnetwork (Figure 2B) were re-analysed using pair-wise domain-class content distance analysis. A similarity threshold of 90% resulted in clade specifc domain-class subnetworks for *Salmonella, Enterobacter* and to a lesser extent for *Klebsiella*. Note that by adopting a whole-genome domainome approach, the history of every domain-class present in the pan-domainome, is taken into account. However, signifying domain classes are the main contributors and similar to what has been observed in Ochman et al.^37^, we observed that the many incidental HGT candidate domain classes appear to have little impact on whole-genome domainome based phylogenetic reconstructions.

The ratio between the core- and pan-domainome size of groups of organisms at different phylogenetic levels provided a good estimate for beta-diversity. A relatively low ratio between the core and pan-domainome reduces the functional assignments that can be inferred from the 16S-rRNA classification. Conversely, a high ratio gives more certainty that functionalities are present. Overall the majority of the analysed families showed a low ratio indicating that only a reduced functional landscape can be extrapolated using 16S-rRNA analysis and the ratio can differ significantly among families. For example, *Chlamydiaceae* shows a large ratio whereas *Enterobacteriaceae* has the lowest observed ratio, indicating that the *Chlamydia* genus which consists mostly of pathogenic bacteria that are obligate intracellular parasites have evolved through simplification instead of complexification and are therefore less diverse^38^. Whereas *Enterobacteriaceae* is a diverse family consisting of members that are part of the gut flora and also contains a wide range of pathogenic species, showing a more diverse functional landscape.

Combining the information from the functional landscape with 16S-rRNA sequences, allowed us to relate the functional diversity with evolutionary distances (Figure 8). This analysis revealed that three stages of diversification can be defined^39^. The first stage represents a fast-short-term evolution, as 16S-rRNA evolutionary distances remain small, though functional diversification has already taken place. This happens in closely related, near identical, related strains where gene acquisition could play a significant role in functional diversity. The second stage represents a long-term evolution, in which functional diversification occurs at a scale compatible with evolutionary time, as reflected by 16S-rRNA evolution. In the third stage diversification of the functional landscape continues but, due to 16S-rRNA genetic constraints, does not align well with 16S-rRNA sequence distances.

## Conclusions

16S-rRNA similarity scores can still be used as a metric for taxonomic classification but we propose a more probalistic interpretation as its performances will be better at higher taxonomic levels.

Whole genome protein domain phylogenies correlate with, and complement 16S-rRNA sequence-based phylogenies. Moreover, domain-based phylogenies reveal rapid functional diversification, allowing for large scale functional comparisons between clades and can be constructed over large evolutionary distances.

Protein domain persistence ratio’s highlight both signifying domain classes and HGT candidates. The increased granularity obtained will pave the way for new applications to better predict the relationships between genotype, physiology and ecology.

## Methods

### Genome annotation

A total of 5713 publicly available complete bacterial genomes were downloaded from the NCBI repository (November 2016)^40^. To prevent technical bias due to the use of different annotation tools and pipelines and different thresholds for assessing the significance of the inferred genetic elements, genomes were consistently structurally and functionally *de-novo* annotated using SAPP^22^, an annotation platform implementing a strictly defined ontology^41^.

16S-rRNA prediction was performed using RNAmmer 1.2^42^. Genes were predicted using Prodigal (2.6.3)^43^ and the identified proteins were functionally annotated using the Pfam library (version 30.0) within InterProScan (version 5.21-60.0)^25,44^. Annotations were automatically converted into RDF according to the GBOL ontology^41^ and loaded into a semantic database for high-throughput annotation and analysis. For the retrieval of information, SPARQL was used (See supplementary file S5 for all queries used).

### Quality analysis

Scaling laws have been identified in the genomic distribution of protein domains^45^. These laws result in linear relationships in the number of domain classes with *n* copies and the total number of domain classes in a genome (See supplementary Figure S5). We have verified the linear relationships in the analysed genomes. These indicators have been used here to further verify the integrity of the assembled genomes^46^. Overall, the previously reported scaling laws also hold true when a higher number of genomes is studied.

### Estimation of pan- and core-domainome size

The estimated number of domain classes in the pan- and core-genomes expected, if the sequences of every existing strain were to be included in the analysis, were computed using binomial mixture models as implemented in the micropan R package^47^ using default values for the parameters. Heap’s analysis as implemented in the micropan R package was used to estimate openness or closeness of the pan-genome using 500 genome permutations and repeating the calculation 10 times.

### Domain persistence

The following formulas were used to calculate persistence ratios

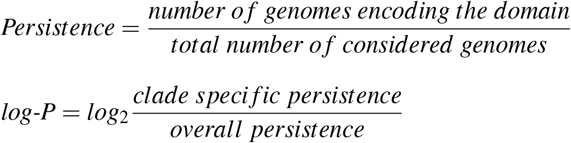

### 16S-rRNA distance calculations

From the *de-novo* annotation, 16S-rRNA sequences were obtained from the semantic database through a SPARQL query (See supplementary file S6 for all queries used). In total 25098 16S-rRNAs were retrieved. rRNA’s that were of low quality (containing N’s) or differed in size greater than the standard deviation were removed from the analysis. Duplicated 16S-rRNAs were merged into a single copy for the multiple alignment. For each 16S-rRNA the orientation was validated using OrientationChecker^48^. The complete gene was used for calculation of pairwise alignment distances using the clustal omega suite for all possible 16S-rRNA pairs (Dataset 1 aligned). The resulting matrix was binarized using 98.7% sequence similarity as a cutoff. The binary matrix was then represented as networks using igraph^49^ in R^50^.

### Domain based distance calculations

Genome distances based on protein domain class content were computed using the asymmetric binary method in which vectors are regarded as binary bits. Non-zero elements are on and zero elements are off. The distance is the proportion of bits in which only one is on amongst those in which at least one is on (dist function in R). A similarity cutoff of ≤ 0.1 was used.

### Statistical software

Statistical analysis and visualisations were performed using R and the following packages, data.table^51^, reshape2^52^, plotly^53^, Biostrings^54^, devtools^55^, micropan^47^, gridExtra^56^, hexbin^57^ and RColorBrewer^58^.

## Supplementary files

All supplementary files can be found at https://doi.org/10.4121/uuid:37346f1e-f8cf-4112-ba4b-1223a3e4edda. This resource contains the entire RDF resource that was used in this study.

## Author’s contributions

JJK, PJS and MSD participated in the conception and design of the study. JJK was responsible for the analysis. JJK, ES, PJS and MSD wrote the manuscript. All authors critically revised the manuscript.

## Acknowledgements

This work was carried out on the Dutch national e-infrastructure with the support of the SURF foundation. This work was partly supported by the European Union’s Horizon 2020 research and innovation programme (EmPowerPutida, Contract No. 635536, granted to Vitor A P Martins dos Santos) and the Netherlands Organisation for Scientific Research funded UNLOCK project (NRGWI.obrug.2018.005) and has received funding form the European Union’s Horizon 2020 research and innovation programme under grant agreement 730976 (IBISBA 1.0).

